# Entropy and codon bias in HIV-1

**DOI:** 10.1101/052274

**Authors:** Aakash Pandey

## Abstract

For the heterologous gene expression systems, the codon bias has to be optimized according to the host for efficient expression. Although DNA viruses show a correlation on codon bias with their hosts, HIV genes show low correlation for both nucleotide composition and codon usage bias with its human host which limits the efficient expression of HIV genes. Despite this variation, HIV is efficient at infecting hosts and multiplying in large number. In this study, first, the degree of codon adaptation is calculated as codon adaptation index (CAI) and compared with the expected threshold value (eCAI) determined from the sequences with the same nucleotide composition as that of the HIV-1 genome. Then, information theoretic analysis of nine genes of HIV-1 based on codon statistics of the HIV-1 genome, individual genes and codon usage of human genes is done. Comparison of codon adaptation indices with their respective threshold values shows that the CAI lies very close to the threshold values. Despite not being well adapted to the codon usage bias of human hosts, it was found that the Shannon entropies of the nine genes based on overall codon statistics of HIV-1 genome are very similar to the entropies calculated from codon usage of human genes. Similarly, for the HIV-1 genome sequence analyzed, the codon statistics of the third reading frame has the highest bias representing minimum entropy and hence the maximum information.

## Introduction

Every organism has its own pattern of codon usage. All the synonymous codons for a particular amino acid are not used equally. Some synonymous codons are highly expressed, whereas the use of others is limited. The use is species-specific [1] [2]. The difference in codon usage is also observed among genes of the same organism [3]. Codon bias has been linked to specific tRNA levels that are mainly determined by the number of tRNA genes that code for a particular tRNA [4]. The choice of codon affects the expression level of genes. This is seen in the expression pattern of transgenes. Gustafsson *et. al.* showed that the use of particular codons can increase expression of the transgene by over 1,000 fold [5]. In bacteria, the gene expressivity correlates with codon usage [6]. Although bacteriophages have been shown to have codons that are preferred by their hosts [7] however, the codon usage pattern of RNA virus differs from its host [8]. Despite this variation, the HIV virus can effectively multiply in human T cells. Codon usage of early genes (*tat, rev, nef*) shows higher correlations with human codon usage [9], but late genes show little correlation. It raises a question how such variation in codon usage still allows for efficient viral gene expression. *van Weringh et. al.* showed that there is a difference in the tRNA pool of HIV-1 infected and uninfected cells. Even though they speculated that HIV-1 modulates the tRNA pool of the host making it suitable for its efficient genome translation, however, the extent to which such modulation helps in efficient translation is still unknown.

After Shannon published his groundbreaking paper “A Mathematical theory of Communication” [10], there have been several attempts in using information theory in the context of living systems. It has been used for measuring the information content of biomolecules, polymorphisms identification, RNA and protein secondary structure prediction, the prediction and analysis of molecular interactions, and drug design [11]. Shannon used the term *information* differently than classical information theorists have used. DNA comprises 4 nucleotides A, G, C, T whose distribution pattern varies among different species. Gatlin deduced information content based on this distribution pattern [12] using transition probability values obtained from the neighbor data [13]. Also, we know that information is not absolute. It depends on the context. This means that the same sequence of DNA may represent different amounts of information depending on what environment it is in or on the machinery that interprets the sequence. We exploit this to calculate the Shannon entropy for the nine genes of HIV-1 based on codon distribution of the viral genome, individual genes and that of its host – human codon usage frequency. Information is calculated based on the codon distribution for three possible reading frames. Here, I have tried to use the information as mentioned by Shannon to see whether information theoretic analysis leads to some novel insights into the problem. To the best of my knowledge, I believe that such study has not been carried out yet. Viruses show overlapping genes and are speculated to be present to increase the density of genetic information [14]. The reason for calculating the Shannon entropy based on three different reading frames is that these genes are read by ribosomal frame-shifting [15]. For those nine genes, I have also calculated the intrinsic entropy of the sequence which can be defined as the entropy based on its own codon usage (i.e. codon usage within the same gene) to compare with other entropy values.

Heterologous expression systems, such as viruses, use host translational machinery for their replication. They are under constant evolutionary pressure to adapt to the host tRNA pool. To estimate a degree of evolutionary adaptiveness of host and viral codon usage, Codon Usage Index (CAI) is used [16]. But, for sequences with a high biased nucleotide composition, interpretation of CAI can be tricky [17]. So, to know whether the value of CAI is statistically significant and has arisen from codon preferences or is merely artifacts of nucleotide composition bias, expected CAI (eCAI) can be the threshold value for comparison [18].

## Methodology

The DNA sequences were obtained from the NCBI database in FASTA format. For each sequence, codon statistics were obtained by entering the sequences on online Sequence Manipulation Suite [19] and by using the standard genetic code as the parameter. Number and fractions of each possible codons were noted. The first nucleotide was deleted to shift the reading frame by +1 to include other possible codon patterns and again the number and frequency were noted. The process was repeated for +2 reading frame. Now, as any of the reading frames can contain the gene of interest, all three reading frame statistics were used to calculate the Shannon entropy separately. The assumption made in the calculation is that reading the message occurs in a linear fashion without slippage of the reading frame (RF).

According to Shannon, for a possible set of events with probability distribution given by {*p1, p2, p3, …, pn*} the entropy or uncertainty is given by,

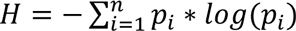

This is, in fact, the observed entropy of a sequence with the given probability distribution. H is the maximum when all *p*_i_ are equally likely. In this condition, the information content is zero. The amount of information or ‘negentropy’ in a sequence can then be given as,

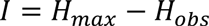

Where, *H*_*obs*_ is the entropy obtained from given probability distribution [20] Lately, the information theoretic value of a given DNA sequence was obtained using the Shannon formula as double sum [21],

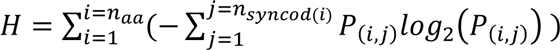

Here, *n*_*aa*_ is the number of distinct amino acids, *n*_*syncod(i)*_ is the number of synonymous codons (or micro-states) for amino acid *i* (or macro-state) whose value range from 1 to 6, and *P*_(*i*,*j*)_ is the probability of synonymous codon *j* for amino acid *i*.

First, a row matrix was constructed with fractions of codons used.

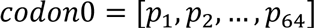

The fractions in the matrix were treated as microstates to calculate Shannon entropy and thus another matrix was constructed consisting of Shannon entropy of each fraction distribution.

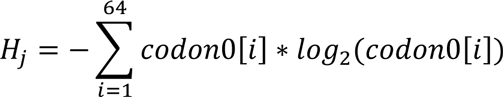

The total Shannon entropy of the sequence is then calculated as:

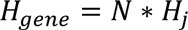

Here *N* is the total number of codons present in the gene of interest. Such calculation was performed for all three RFs. As null models, two nucleotides compositions one with the same nucleotides composition as that of the HIV-1 genome sequence and another with the equal percentages of each four nucleotides (25% each) were used for the construction of random sequences.

The correlation coefficient for each gene’s codon statistics and human codon usage statistics was calculated. Correlation coefficients for two genes *vpr* and *vpu* were calculated again removing the codon data for which no amino acid is present in that gene. Then, Shannon entropy was calculated for all nine genes using the human codon usage statistics. Intrinsic entropy, which is the entropy based on own codon statistics of each gene was also calculated. Again, the assumption is that there is no slippage of reading frame during translation of the message. Thus, codon statistics for single reading frame starting with start codon was used to calculate intrinsic entropy. Similarly, average entropy was calculated by averaging the fractions of codons for all three RFs. For the calculation of percentage overall GC content and position specific GC content of codons of nine genes and CAI values, http://genomes.urv.cat/CAIcal/ [21] online site was used again using the standard genetic code as the parameter. For calculation of the expected codon adaptation index was performed in E-CAI server (http://genomes.urv.es/CAIcal/) using Markov chain and standard genetic code as the parameters. Human codon usage statistics were obtained from the online site (http://genomes.urv.cat/CAIcal/CU_human_nature.html). Computations were performed in R.

## Result and Discussion

The calculation of CAI shows that all the genes have high values (Table 1) albeit with a varying degree of GC content. All the CAI values are greater than 0.6 with *vpu* having the lowest value of 0.62 whereas *tat* has the highest value of 0.77. CAI well above 0.5 is usually considered to be showing a good level of adaptation towards the host, however, one should be careful while interpreting these values as they may not reveal the level of adaptation just by themselves. Such values may result due to the bias in nucleotides composition. So, to know whether these values actually represent the adaptation we need to set a threshold set by the bias in nucleotides composition. For that expected CAI (eCAI) is calculated and compared with CAI [18]. From these comparisons, none of the genes seem to be well adapted to the human codons usage pattern. However, they do not show poor adaptation either as all genes have higher CAI values. Interestingly, they lie close to the threshold values. We can note that the GC content of all the genes is below average. Also, GC content of second and third nucleotides of codons shows the greatest variability. As neither nucleotide bias nor the pressure for adaptation exclusively explains high CAI values, we can speculate that both factors play a role, to some extent, in determining the values.

**Table 1:** Codon adaptation index (CAI) and GC content of HIV-1 genes.

| Gene | Length | CAI | %GC | %GC1 | %GC2 | %GC3 | eCAI (p<0.05) |
| --- | --- | --- | --- | --- | --- | --- | --- |
| *gag* | 1503 | 0.73 | 44.0 | 50.3 | 43.5 | 38.3 | 0.74 |
| *nef* | 621 | 0.76 | 49.4 | 58.0 | 44.4 | 45.9 | 0.76 |
| *tat* | 2592 | 0.77 | 39.8 | 38.8 | 41.0 | 39.7 | 0.78 |
| *pol* | 1746 | 0.71 | 38.3 | 48.8 | 36.4 | 29.6 | 0.72 |
| *rev* | 2682 | 0.73 | 40.5 | 41.5 | 39.7 | 40.2 | 0.74 |
| *vif* | 579 | 0.72 | 42.0 | 46.6 | 41.5 | 37.8 | 0.74 |
| *vpr* | 291 | 0.73 | 45.0 | 54.6 | 34.0 | 46.4 | 0.74 |
| *vpu* | 249 | 0.62 | 37.8 | 54.2 | 31.3 | 27.7 | 0.66 |

CAI and eCAI values with overall GC percentage and position specific GC percentage for nine HIV-1 genes is reported. %GC 1, %GC2 and %GC3 represent GC percentage of the first, second and the third position of the codons respectively. Second and third position of the codon shows greatest bias in GC content as compared to the first position (except for *tat* and *rev* gene which shows almost no bias for all three positions)

Entropic calculation shows a general trend for the sequences analyzed. First, +2 frame-shifted reading frame shows lower entropy as compared to two other reading frames. This marked distinction of the third reading frame among three possible reading frames of the sequence analyzed is surprising. As it has the lowest entropy among three reading frames, the sequence with the codon usage pattern of the third RF represents the highest information (in Shannon’s sense) (Table 3). Calculations from 10 random genomic sequences, having the same nucleotide bias as that of the HIV-1 genome, did not show such pattern. The mean of the average entropy per codon for each reading frames of those random sequences was 5.8 with the standard deviation of 0.05. This probably suggests that there is a genome-wide conservation of codon usage for the third reading frame of the HIV genome analyzed, but the reason is unclear.

We see that there is a high correlation between codon present in HIV-1 early genes (*tat, rev, nef*) and human codon usage (fig 1), but correlation is lower for other genes: *env, gag, pol, vpr, vif, vpr* (table 2); *vpu* and *vif* genes have the lowest correlation with human codon usage. The degree of correlation also differs for 3 RFs’ codon statistics with that of nine genes: high correlation probably suggesting that the particular gene resides at that RF.

**Fig 1:**
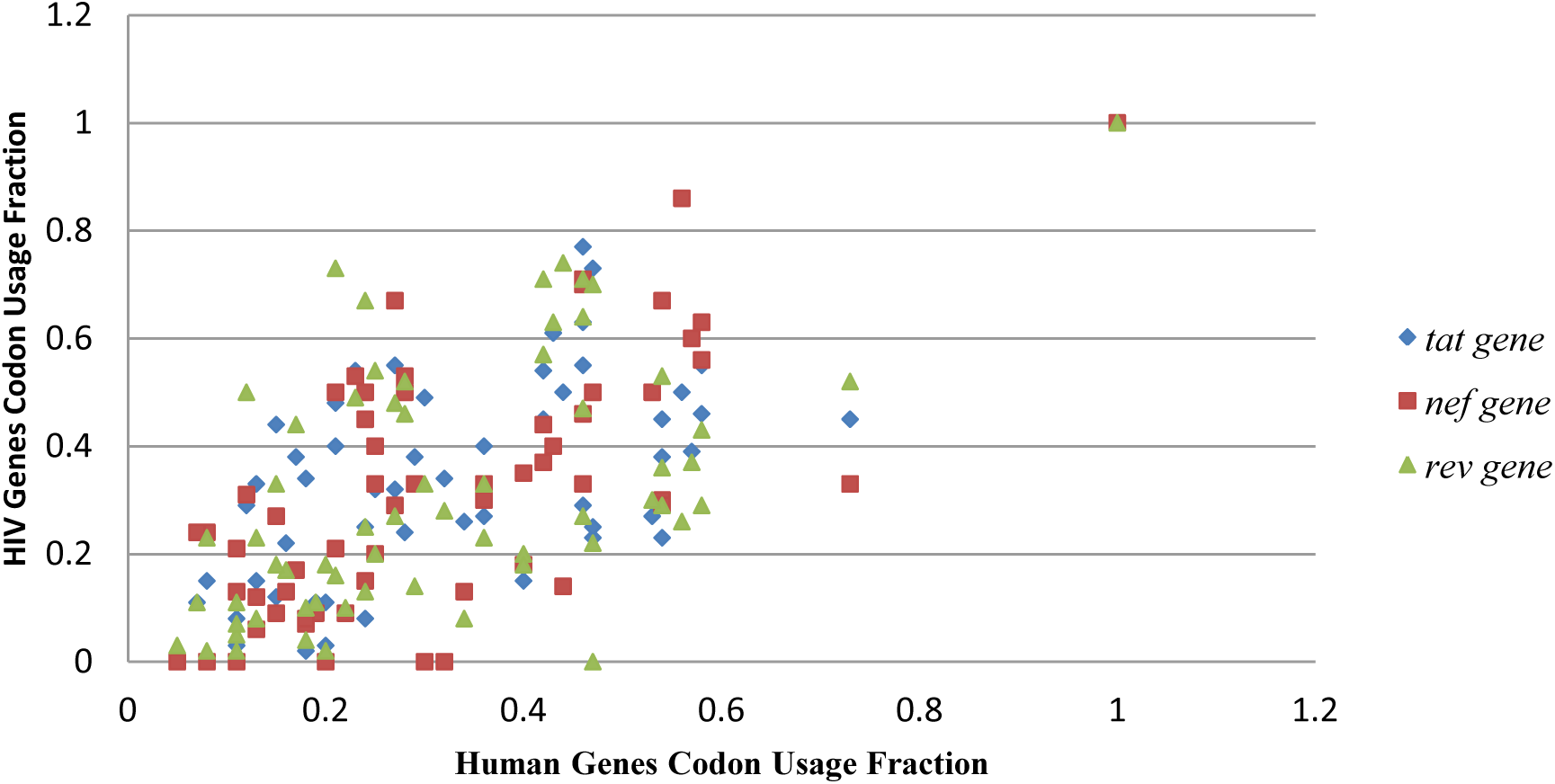
Scatter plot between codons fractions of *tat, rev* and *nef* genes of HIV-1 and human codon usage fraction with the correlation coefficient of 0.72, 0.62 and 0.73 respectively. These three genes have the highest correlation with human codon usage and are also the early genes.

**Table 2.** Correlation coefficients calculated among human codon usage, codon usage of HIV-1 genes and codon statistics for all three reading frames of HIV-1 genome.

| Genes | Correlation coefficients |  |  |  |
| --- | --- | --- | --- | --- |
|  | Human codon | RF 0 | RF +1 | RF +2 |
| *env* | 0.48 | 0.78 | 0.74 | 0.87 |
| *gag* | 0.55 | 0.84 | 0.76 | 0.83 |
| *tat* | 0.72 | 0.88 | 0.96 | 0.91 |
| *rev* | 0.62 | 0.91 | 0.87 | 0.94 |
| *vpu* | 0.26 / 0.41 | 0.61 | 0.54 | 0.68 |
| *vpr* | 0.48 / 0.55 | 0.61 | 0.66 | 0.65 |
| *vif* | 0.44 | 0.79 | 0.74 | 0.76 |
| *nef* | 0.73 | 0.78 | 0.76 | 0.73 |
| *pol* | 0.57 | 0.84 | 0.82 | 0.89 |
| **Human** | - | 0.74 | 0.78 | 0.66 |

The table shows the correlation between codon fractions of nine genes with human codon usage fraction (column 2) and also with three different reading frames. RF 0 represents the codon fractions of the initial reading frame of HIV-1 sequence, RF +1 represents a codon fractions of single frame shift and RF +2 represents codon fractions of +2 frameshifts of the HIV-1 genome sequence. Human codon usage statistics is also compared with the three possible reading frames of HIV-1 genome where +1 shifted reading frame shows the highest correlation. For *vpu* and *vpr* gene, the number after backlash is calculated removing the data for which no codons are present for certain amino acids.

*env, rev, pol* and *vpu* genes have the highest correlation with the third reading frame as compared to other two reading frames. Similarly, *gag* and *vpr* also have a high correlation coefficient. If we use codon statistics of third RF to calculate the Shannon entropy, we get the minimum entropy and hence maximum information. But, then again we run into a problem as this third RFs shows the lowest correlation coefficient with human codon usage pattern. So, there must be a balance between these contrasts: maximizing information or maximizing correlation. Take *gag* for example, it shows high correlation with RF0 and RF2 both of which have lower correlation with human codon usage. This means that the choice of codons affects the *gag* gene expression. In fact, the ratio of native and optimized codons determines the HIV-1 *gag* expression [23]. This also supports the speculation that codon bias leads to sub-optimal expression in infected cells. There is, in fact, good evidence that HIV-1 gene expression is not the maximum but, is finetuned to allow regulation of diverse processes [24]. More evidence of sub-optimal expression is shown by the fact that when codon optimized genes that are better adapted to the host tRNA pool were introduced, it led to higher expression [25][26][27]. From the entropic calculation based on human codon usage, we can see that *vpu* and *vif* have the lowest entropy, but as they have a low correlation with human codon usage, their expression is limited. Codon optimization of these genes results in the increase of expression level [28]. However, high correlation does not imply that the gene is in that reading frame. It is possible that such bias may or may not affect biological function, but it is likely that such distinction of lower entropy has some evolutionary importance.

**Table 3:**
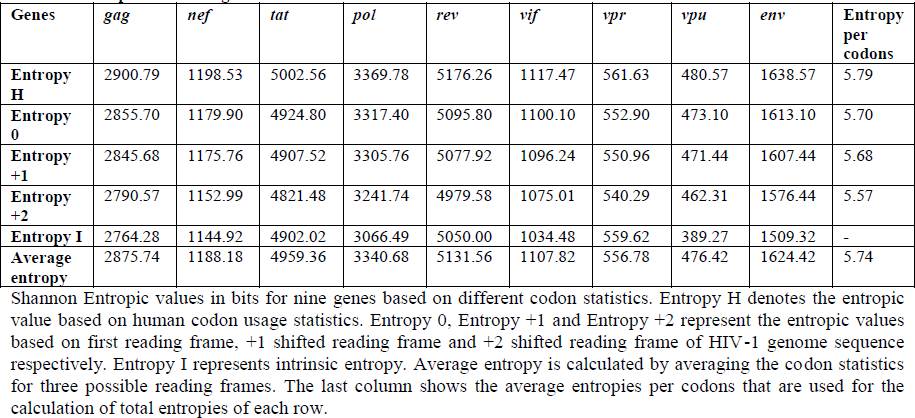
Entropies of HIV-1 genes based on various codon distributions.

| Genes | *gag* | *nef* | *tat* | *pol* | *rev* | *vif* | *vpr* | *vpu* | *env* | Entropy per codons |
| --- | --- | --- | --- | --- | --- | --- | --- | --- | --- | --- |
| Entropy H | 2900.79 | 1198.53 | 5002.56 | 3369.78 | 5176.26 | 1117.47 | 561.63 | 480.57 | 1638.57 | 5.79 |
| Entropy 0 | 2855.70 | 1179.90 | 4924.80 | 3317.40 | 5095.80 | 1100.10 | 552.90 | 473.10 | 1613.10 | 5.70 |
| Entropy +1 | 2845.68 | 1175.76 | 4907.52 | 3305.76 | 5077.92 | 1096.24 | 550.96 | 471.44 | 1607.44 | 5.68 |
| Entropy +2 | 2790.57 | 1152.99 | 4821.48 | 3241.74 | 4979.58 | 1075.01 | 540.29 | 462.31 | 1576.44 | 5.57 |
| Entropy I | 2764.28 | 1144.92 | 4902.02 | 3066.49 | 5050.00 | 1034.48 | 559.62 | 389.27 | 1509.32 | - |
| Average entropy | 2875.74 | 1188.18 | 4959.36 | 3340.68 | 5131.56 | 1107.82 | 556.78 | 476.42 | 1624.42 | 5.74 |

Shannon Entropic values in bits for nine genes based on different codon statistics. Entropy H denotes the entropic value based on human codon usage statistics. Entropy 0, Entropy +1 and Entropy +2 represent the entropic values based on first reading frame, +1 shifted reading frame and +2 shifted reading frame of HIV-1 genome sequence respectively. Entropy I represents intrinsic entropy. Average entropy is calculated by averaging the codon statistics for three possible reading frames. The last column shows the average entropies per codons that are used for the calculation of total entropies of each row.

Intrinsic entropy differs greatly with other entropic values as it shows lowest values for most genes. Such low intrinsic entropies may have significance for free-living organisms as lower entropies suggest higher bias. But, for heterologous expression systems such as HIV-1, entropy H probably represents the best entropic values for the genes analyzed as host (human) gene usage codon statistics was used for the calculation. Average entropy, which is closer to entropy H rather than intrinsic entropy, gives a better representation for entropic value and hence for the amount of information a gene contains inside a human host. Although there is great variation in the synonymous codon usage statistics between HIV-1 genes and human genes, the entropic values for the HIV-1 genes based on the overall code distribution of the HIV genome shows almost similar values as compared to the calculation based on human codon usage statistics (Table 3). Even for *a vpu* gene, which has a very low correlation coefficient (0.26), the entropic values based on overall codon statistics of HIV-1 genome and human codon usage statistics show similarity: 476.42 and 480.57 bits respectively. Even if we remove the data for which there is no single codon for certain amino acids in that gene, the correlation coefficient is still low. In *vpu* gene, codons for Cysteine, Threonine and Phenylalanine are absent. If we remove that data, we get a correlation coefficient of 0.41, which is still low. However, this removal does not affect the entropic calculation. Similarly, for *vpr* gene, codons for Cysteine are absent. Removing that data new correlation coefficient obtained is 0.55 and average entropic values and entropy H are close: 556.78 and 561.63 bits respectively.

HIV is a highly variable virus which undergoes rapid mutation. Although it cannot match its codon bias with that of the host, but it can have a stable codon usage pattern. To maintain the overall codon statistics, it has to maintain the nucleotide composition, which is the determinant of codon bias [28]. Despite its high mutation rate, the biased nucleotide composition of HIV is constant over time [29]. If the genes have same codon biases as that of the host then it might lead to their highest expression. But this is not desired as it would not allow for efficient tuning of its complex processes. If codon bias is completely different from that of the host, then it might result in very low expression putting its ability to survive in the host in question. So, HIV has to find a solution which can result in sub-optimal expression of the genes. From the calculation in table 3 (Entropy H and Average Entropy), it seems that HIV has found a solution in which its codon bias is different from that of host to allow sub-optimal expression, but at the same time represent the same level of information as can be obtained from the codon bias of its host. Also, random sequences generated from the conserved nucleotide bias (but not from random sequences with no nucleotide bias) give the same results suggesting that nucleotide bias decides the entropic values (Table 4). In fact, both average entropies per codon calculated from HIV-1 genome and the human genes (Table 3) lie within 95% confidence interval of the mean of the average entropy per codon obtained from random sequences generated from the conserved nucleotide bias. One possible explanation for this observation is that, despite having a difference in codon bias with human, HIV-1 viruses have evolved to represent the same level of information as would have represented by the codon bias of the human host.

**Table 4:** Entropic calculation and ANOVA for random sequences.

|  | Mean of average entropy per codon | 95% of confidence interval | Standard Deviation | F-value | Prob>F |
| --- | --- | --- | --- | --- | --- |
| RC Sequences | 5.76 | 5.72-5.80 | 0.04 | 136.23 | 7.87E-10 |
| RNC Sequences | 5.97 | 5.95-5.97 | 0.02 |  |  |

Mean average entropy per codons for random sequences and ANOVA for the result obtained is reported. RC represents ten random sequences generated from conserved nucleotide distribution. This is the same nucleotide distribution as that of the HIV-1 genome. RNC represents another ten random sequences that are generated with equal fractions of nucleotides (0.25 each). ANOVA shows that the F-value is large 136.23 with very low p>F suggesting the means of entropies from the two distributions are different.

## Conclusion

Despite many studies, HIV viral genome still possesses several mysteries. HIV is evolving along with its human host. However, it is not clear why its nucleotide composition and synonymous codon usage bias differ greatly from its host. From the comparison of CAI with eCAI, we can conclude that HIV genes are not well adapted to the tRNA pools of humans. So, it can be inferred that selection pressure on HIV to adapt to tRNA pools is counteracted by the rapid mutation of its genome. It is not clear whether nucleotide composition bias can give rise to the asymmetry in the observed information content along three possible reading frames. However, despite having large differences in nucleotide composition and synonymous codon usage bias, HIV genes are seem to have evolved to represent the same level of information as obtained by the codon bias of human genes. How HIV is able to attain such uniformity, despite differing from its host, is yet another mystery this study has surfaced. Further work is needed, which can bring together the differences in one place to give a clear picture of the evolution of the HIV viral genome.

